# A high-quality chromosome-level genome of an undescribed *Meretrix* species using Nanopore and Hi-C technologies

**DOI:** 10.1101/2025.04.01.646383

**Authors:** Che-Chun Chen, Te-Hua Hsu, Hsin-Yun Lu, Sen-Lin Tang, Ying-Ning Ho

**Affiliations:** Biodiversity Research Center, Academia Sinica, Taipei, Taiwan; Taiwan Oceans Genome Center, National Taiwan Ocean University, Keelung, Taiwan; Department of Aquaculture, National Taiwan Ocean University, Keelung, Taiwan; Institute of Marine Biology, National Taiwan Ocean University, Keelung, Taiwan; Center of Excellence for the Oceans, National Taiwan Ocean University, Keelung, Taiwan

**Author notes:** Correspondence: Ying-Ning Ho.

## Abstract

*Meretrix* is a commercially valuable bivalve genus in Asia, but only one reference genome has hindered comprehensive genetic studies and germplasm resource evaluation. In this study, we present three reference genomes of *Meretrix* species: *Meretrix* sp. MF1, *Meretrix* sp. MT1, and *Meretrix lamarckii* JML1. *Meretrix* sp. MF1 was assembled at the chromosome level using Nanopore sequencing and Hi-C technologies, whereas *Meretrix* sp. MT1 and *Meretrix lamarckii* were assembled as scaffold-level assemblies. The chromosome-level genome of *Meretrix* sp. MF1 consists of 36 contigs, including 19 chromosomes and 17 scaffolds, with a total length of 883.3 Mb and a scaffold N50 of 46.87 Mb. Notably, the genome of *Meretrix* sp. MF1, a putative novel species, exhibits an Average Nucleotide Identity (ANI) of 94.33% with its closest relative, *Meretrix lamarckii*. These genomic resources not only provide a crucial foundation for genetic research on Meretrix but also contribute to the development of effective conservation strategies for its sustainable management.

## Background & Summary

The genus *Meretrix* is a commercially significant marine bivalve widely distributed across the warm coastal waters of East and Southeast Asia^1^. It is particularly abundant along the southern Taiwan coastline, where it has become one of the most economically valuable species in aquaculture^2^. *Meretrix* thrives in water temperatures ranging from 25°C to 33°C, with significant growth slowing below 20°C and mass mortality occurring when temperatures exceed 45°C. Additionally, it prefers salinities between 16 and 35 ppt, with extreme fluctuations in salinity adversely affecting its survival and development^3^. In recent years, large-scale mortality events and slow growth have posed a significant threat to *Meretrix* aquaculture in Taiwan. Previously, a single hectare of culture area could yield up to 18 metric tons, but current yields have plummeted to as low as 0.6 metric tons^4^. Several factors are thought to contribute to this decline, including environmental degradation, climate change, disease outbreaks, improper aquaculture management, and genetic deterioration due to inbreeding^5^.

Despite the economic and ecological significance of *Meretrix*, genomic resources for this genus remain scarce. To date, the genome of only *M. petechialis* has been published (GCA_046203225.1), and the morphological similarities among various *Meretrix* species present challenges for accurate classification and genetic studies. A high-quality reference genome is essential for understanding the genetic basis of adaptive evolution, population dynamics, and potential genetic vulnerabilities within *Meretrix* species. Moreover, genomic data could shed light on mechanisms underlying disease resistance, stress tolerance, and reproductive strategies, all of which are critical for the sustainable management and conservation of these species. *Meretrix* species are commonly found in the coastal and estuarine areas of Taiwan. However, these two habitats exhibit distinct environmental conditions. Coastal waters typically maintain higher salinity levels, ranging from 32 to 35 psu, whereas estuarine areas experience greater salinity fluctuations, potentially varying from 0.5 to 35 psu. Therefore, in this study, we collected *Meretrix* samples from these two contrasting environments. *Meretrix* sp. MF1 was specifically collected from the open coastal waters (Anping, Tainan), while *Meretrix* sp. MT1 was exclusively obtained from the estuarine environment (Cigu, Tainan). Our Previously study has showed that the *Meretrix lamarckii* clade is divided into two main distinct groups: one containing sample collect from Japan, and the other containing samples from Taiwan, suggesting that *M. lamarckii* from Taiwan and *M. lamarckii* from Japan are distinct species^6^. Therefore, we selected *Meretrix* sp. MF1, a potential novel species, for high-quality chromosome-level genome assembly. As there is no reference genome for *M. lamarckii* currently, *M. lamarckii* JML1 from Japan was also selected for genome assembly. On the other hand, *Meretrix* sp. MT1, MT2, and MT3, collected from the coastal waters of Taiwan, formed a distinct clade and were most closely related to *M. lusoria* from China. *Meretrix* sp. MT1 was selected for genome assembly.

In this study, we present chromosome-level genome assemblies of one *Meretrix* species, using a combination of Illumina short-read sequencing, Nanopore long-read sequencing, and Hi-C chromatin conformation capture technologies. For *Meretrix* sp. MF1, we generated a total of 51.1 Gb of Illumina data, 80.02 Gb of Nanopore data, and 46.48 Gb of Hi-C data. The final assembly yielded 19 chromosomes with a total length of approximately 883.3 Mb and a scaffold N50 of 46.87 Mb. Based on this high-quality reference genome, we successfully assembled the genomes of two additional *Meretrix* species, *Meretrix* sp. MT1 and *M. lamarckii* JML1. For *Meretrix* sp. MT1, we obtained 56.6 Gb of Illumina data and 66.79 Gb of Nanopore data, resulting in the assembly of 19 chromosomes with a total length of 944.74 Mb. Similarly, for *M. lamarckii* JML1, we obtained 42.6 Gb of Illumina data and 88.91 Gb of Nanopore data, resulting in the assembly of 19 chromosomes with a total length of 883.07 Mb. To further explore the genetic relationships among these species, we conducted comparative genomic analyses, including nuclear genome alignment, mitochondrial genome comparisons, and average nucleotide identity (ANI) calculations. Our results revealed that *Meretrix* sp. MF1 and *M. lamarckii* JML1 exhibit the highest similarity, with an ANI of 94.33%. This genomic divergence suggests that *Meretrix* sp. MF1 represents a novel *Meretrix* species. Based on these findings, we propose the designation *M. formosana* for this newly identified species.

The high-quality reference genome presented in this study provides a valuable foundation for future research on *Meretrix* population genomics, adaptive evolution, and genetic diversity. It will also facilitate further studies on gene function, aquaculture enhancement, and sustainable aquaculture practices. Additionally, our findings highlight the importance of genomic resources in identifying cryptic species, understanding evolutionary processes, and supporting sustainable aquaculture efforts. The availability of this genomic data will empower researchers and aquaculture practitioners to develop targeted breeding programs and genetic management strategies, ultimately enhancing the resilience and productivity of *Meretrix* populations in the face of environmental challenges.

## Methods

### Sampling and nucleic acid extraction

Samples of *Meretrix* sp. MF1 were collected from the coastal waters of southern Taiwan (Anping, Tainan), while *Meretrix* sp. MT1 was obtained from the estuarine region of southern Taiwan (Cigu, Tainan). *M. lamarckii* JML1 was sourced from GOURMET HUNTER CO., LTD., a Taiwan-based international trading company specializing in aquatic products. The sample originated from Chiba, Japan. Genomic DNA was extracted from 25 mg of muscle tissue using the Nanobind^®^ PanDNA Kit (PacBio, USA) following the ‘Extracting DNA from animal tissue using the Nanobind^®^ PanDNA kit’ protocol. The extracted DNA was stored at -80°C to preserve its integrity. DNA quality was assessed using 1.0% agarose gel electrophoresis, fluorescence quantification with the Qubit™ 4 Fluorometer (Thermo Fisher Scientific, USA) with Qubit™ dsDNA BR Assay Kits (Thermo Fisher Scientific, USA), as well as spectrophotometric analysis using the NanoDrop™ One Microvolume UV-Vis Spectrophotometer (Thermo Fisher Scientific, USA).

### Phylogenetic analysis of *Meretrix* species

There are 33 Cytochrome c oxidase subunit I (COXI) sequences from *Meretrix* species were selected for phylogenetic analysis, 27 sequences from NCBI database (*M. lamarckii, M. lusoria, M. lyrate, M. meretrix*, and *M. petechialis*) and six from this study (*M. lamarckii* JML1, JML2 and *Meretrix* sp. MF1, MT1, MT2, MT3). A neighbor-joining tree was constructed using MEGA version 11.0.13^7^, with 1000 bootstrap replicates and the Tamura-Nei model.

### Library preparation and sequencing

Genomic DNA was purified using AMPure XP Reagent (Beckman Coulter, USA) following the manufacturer’s protocol, and each purified sample was quantified using the Qubit™ 4 Fluorometer with Qubit™ dsDNA BR Assay Kits. Nanopore sequencing libraries were prepared using SQK-LSK110 Ligation Sequencing Kit (Oxford Nanopore Technologies, UK) according to the manufacturer’s protocol. A 150 µL aliquot of the library was loaded onto FLO-PRO002 (R9.4.1) flow cells (Oxford Nanopore Technologies, UK) for the PromethION 2 Solo (Oxford Nanopore Technologies, UK), and sequenced for approximately 120 hrs. The reads were then basecalled using Dorado version 0.7.0 (https://github.com/nanoporetech/dorado) with the super-accurate (SUP) model, yielding 80.02 Gb of data with 6.76 M high-quality reads for *Meretrix* sp. MF1 (Table 1). Additionally, the data for *Meretrix* sp. MT1 and *M. lamarckii* JML1 are summarized in Table 1. Illumina sequencing libraries were constructed using the TruSeq^®^ Nano DNA Library Prep Kit (Illumina, USA) following the manufacturer’s guidelines. Genomic DNA was fragmented to approximately 350 bp via sonication, purified with Sample Purification Beads (Illumina, USA), and sequenced on the NovaSeq X Plus System (Illumina, USA), producing 150 bp paired-end reads. The raw Illumina reads, averaging 50.1 Gb per sample, were processed using fastp version 0.23.4^8^ for quality control (Table 1). For chromosome-level assembly, the Hi-C library was constructed using the Dovetail^®^ Omni-C^®^ Kit (Cantata Bio, USA) following the manufacturer’s protocol. The library quality was assessed using a Qsep 100 Bio-Fragment Analyzer (BiOptic, Taiwan) with an S2 Standard Cartridge Kit (BiOptic, Taiwan) and a Qubit™ 4 Fluorometer with Qubit™ dsDNA HS Assay Kits (Thermo Fisher Scientific, USA). The library was then sequenced on the Novaseq X Plus System, generating 150 bp paired-end reads and yielding 46.48 Gb of data, with 309.86 M reads (Table 1).

**Table 1.** Statistics for the sequencing data of the *Meretrix* genome.

| Species | Platform | Reads (M) | Raw Data (Gb) | Average Read Length (bp) | Maxmiun Length (bp) | N50 Read Length (bp) | Coverage (X) |
| --- | --- | --- | --- | --- | --- | --- | --- |
| *Meretrix* sp. MF1 | Illumina | 340.85 | 51.10 | 150 | 150 | 150 | 57.69 |
|  | Nanopore | 6.76 | 80.02 | 11,830 | 1,967,475 | 26,493 | 90.33 |
|  | Hi-C | 309.86 | 46.48 | 150 | 150 | 150 | 52.47 |
|  | Total | - | 177.60 | - | - | - | 200.49 |
| *Meretrix* sp. MT1 | Illumina | 377.55 | 56.60 | 150 | 150 | 150 | 60.01 |
|  | Nanopore | 20.64 | 66.79 | 3,236 | 1,850,522 | 5,915 | 70.81 |
|  | Total | - | 123.39 | - | - | - | 130.82 |
| *M. lamarckii* JML1 | Illumina | 283.81 | 42.60 | 150 | 150 | 150 | 45.97 |
|  | Nanopore | 71.80 | 88.91 | 1,238 | 2,205,180 | 2,159 | 95.95 |
|  | Total | - | 131.51 | - | - | - | 141.92 |

### Genome assembly and scaffolding

The general workflow of this study is illustrated in Fig. 1. Draft genome for *Meretrix* sp. MF1 and *Meretrix* sp. MT1 were generated using Nanopore data processed with Nextdenovo version 2.5.2^9^. However, due to the shorter read lengths in *M. lamarckii* JML1 Nanopore data, its genome was assembled using Masurca version 4.1.2^10^. The data were then processed with NanoFilt version 2.8.0^11^ with Q12 for quality control. Next, both Nanopore and Illumina data were integrated and polished with Nextpolish version 1.4.1^12^ followed by Purge_Dups version 1.2.6^13^ to remove redundant sequences. Hi-C data was utilized to construct the chromosome-level genome assembly for *Meretrix* sp. MF1. Initially, fastp version 0.23.4^8^ was employed for quality control, and Chromap version 0.2.7^14^ was used for alignment and preprocessing. Scaffolding was carried out using YaHS version 1.2.2^15^ to generate chromosome-level scaffolds. Subsequently, Juicer tools version 2.20.00^16^ was applied to construct the Hi-C contact matrix and contact map. The resulting chromosome-level genome assembly for *Meretrix* sp. MF1 had a total length of 883.3 Mb, with a longest scaffold of 59.29 Mb, an N50 of 46.87 Mb, and an L90 of 17 (Table 2). The Hi-C map (Fig. 2A) revealed 19 chromosome-scale scaffolds, which collectively accounted for 99.54% of the total genome size. Chromosome sizes ranged from 28.62 Mb to 59.29 Mb, with an average length of 46.27 Mb (Table 3). The genome was further visualized using TBtools-II version 2.156^17^ (Fig. 2B). To refine and scaffold the genomes of *Meretrix* sp. MT1 and *M. lamarckii* JML1, RAGTAG version 2.1.0^18^ was used, with *M. petechialis* (GCA_046203225.1) serving as the reference genome for *Meretrix* sp. MT1, and *Meretrix* sp. MF1 as the reference for *M. lamarckii* JML1. Redundant sequences were then filtered using Purge_Dups version 1.2.6^13^, and Nextpolish version 1.4.1^12^ was applied for a final round of genome refinement. The final assembly details for all three species are summarized in Table 3.

**Table 2.** The assembly statistics of *Meretrix* genome.

| Species | *Meretrix* sp. MF1 | *Meretrix* sp. MT1 | *M. lamarckii* JML1 |
| --- | --- | --- | --- |
| BioSample Accession | SAMN47132296 | SAMN47132297 | SAMN47132298 |
| Assembly level | Chromosome | Scaffold | Scaffold |
| Contig/Scaffold | 36 | 74 | 198 |
| N50 | 46,874,007 | 48,788,946 | 46,538,327 |
| N90 | 39,672,059 | 39,783,319 | 39,783,319 |
| L50 | 9 | 9 | 9 |
| L90 | 17 | 17 | 17 |
| Total length | 883,299,404 | 944,737,021 | 883,065,649 |
| Maximum length | 59,289,571 | 61,949,085 | 61,949,085 |
| Mean length | 24,536,094.56 | 12,766,716.50 | 4,459,927.52 |

**Table 3.** The 19 chromosomes length (bp) of *Meretrix* genome.

| Chromosome | *Meretrix* sp. MF1 | *Meretrix* sp. MT1 | *M. lamarckii* JML1 |
| --- | --- | --- | --- |
| Chr1 | 59,289,571 | 61,949,085 | 59,194,310 |
| Chr2 | 54,965,621 | 60,624,939 | 55,663,535 |
| Chr3 | 53,900,049 | 59,811,477 | 55,047,926 |
| Chr4 | 52,859,844 | 58,379,201 | 52,423,577 |
| Chr5 | 52,363,401 | 56,111,155 | 52,249,500 |
| Chr6 | 50,989,597 | 56,103,256 | 51,210,398 |
| Chr7 | 50,412,902 | 55,667,226 | 50,132,483 |
| Chr8 | 48,000,840 | 52,361,717 | 50,102,710 |
| Chr9 | 46,874,007 | 48,788,946 | 46,538,327 |
| Chr10 | 45,839,534 | 48,684,485 | 46,236,359 |
| Chr11 | 45,293,453 | 47,168,644 | 44,711,411 |
| Chr12 | 44,574,984 | 46,261,984 | 42,644,817 |
| Chr13 | 44,295,794 | 45,430,347 | 41,753,149 |
| Chr14 | 43,353,122 | 44,783,010 | 41,397,296 |
| Chr15 | 43,345,231 | 43,390,135 | 41,233,522 |
| Chr16 | 40,473,790 | 42,721,536 | 40,946,197 |
| Chr17 | 39,672,059 | 41,343,720 | 39,783,319 |
| Chr18 | 34,081,375 | 35,115,754 | 32,822,444 |
| Chr19 | 28,621,390 | 33,208,588 | 27,664,891 |
| Mean | 46,274,029.68 | 49,363,431.84 | 45,881,903.74 |
| Total | 879,206,564 | 937,905,205 | 871,756,171 |
| Percentage (%) | 99.54 | 99.28 | 98.72 |
| Unplaced | 4,092,840 | 6,831,816 | 11,309,478 |

**Figure 1.**
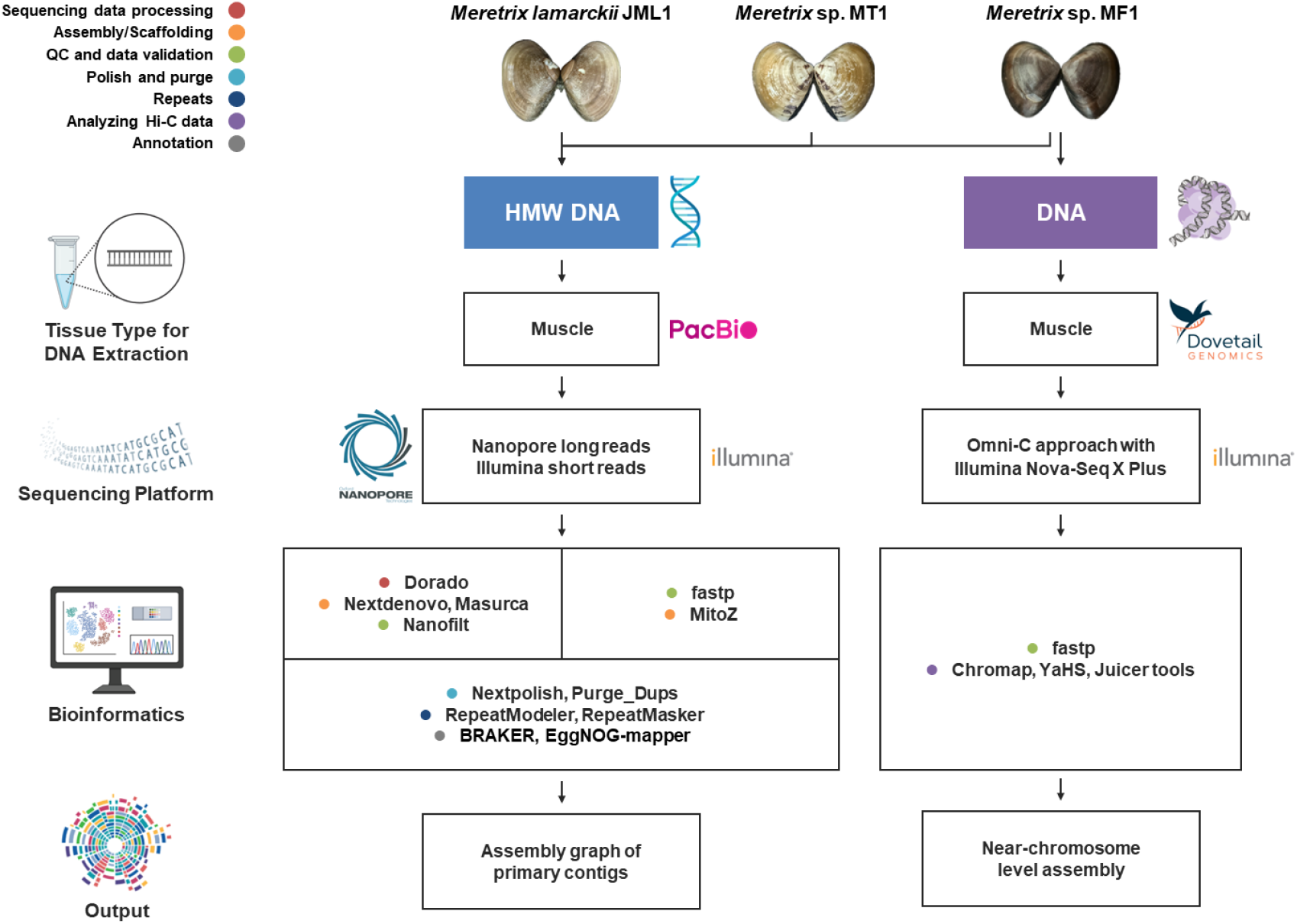
Schematic overview of the general workflow.

**Figure 2.**
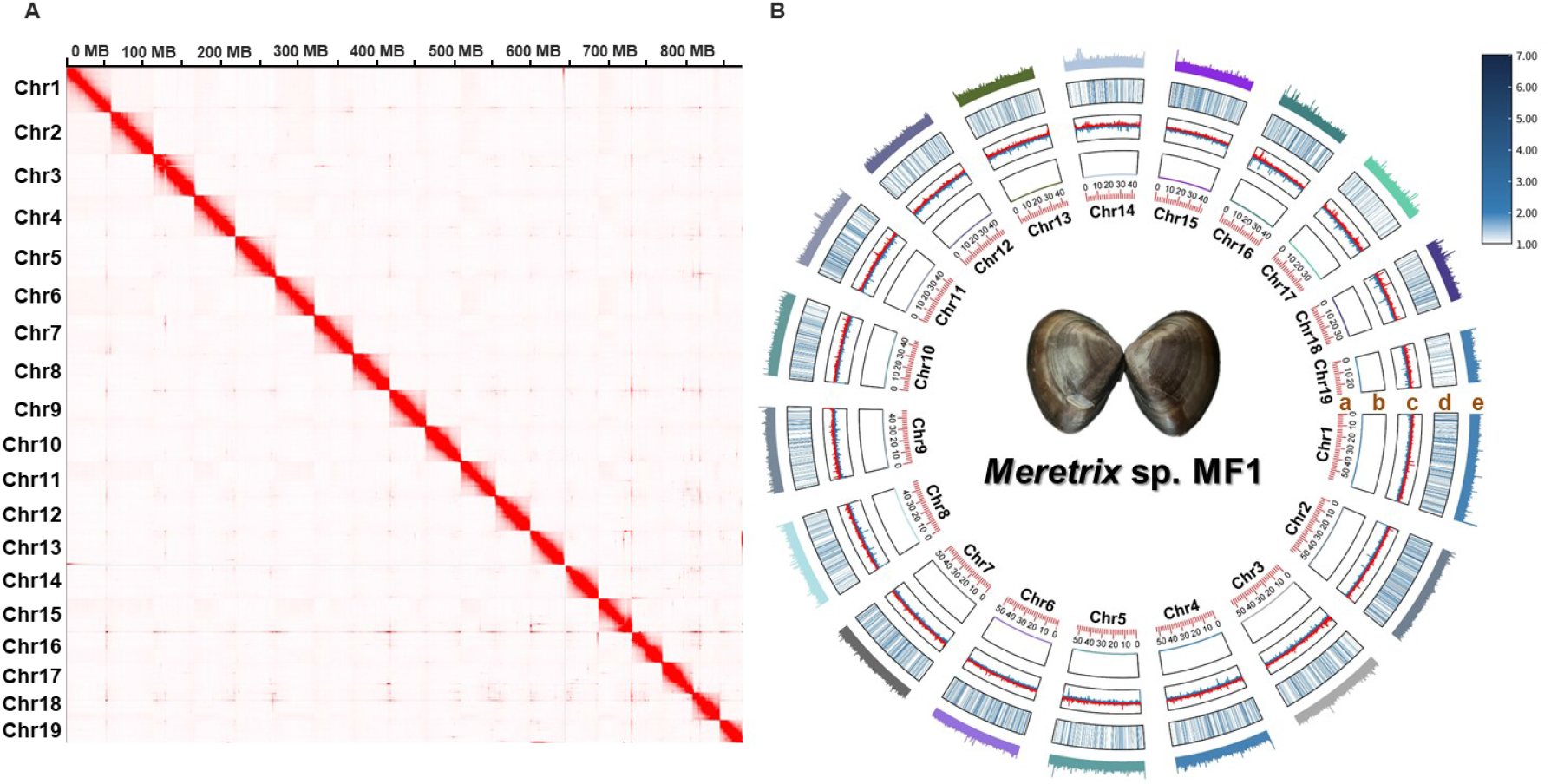
Characteristics of *Meretrix* sp. MF1 genome assembly. (**A**) Hi-C heatmap of chromosomal interactions in the *Meretrix* sp. MF1 genome. (**B**) A circos plot of the *Meretrix* sp. MF1 genome, with tracks from innermost to outermost as follows: (a) Numbers and sizes of *Meretrix* sp. MF1 chromosomes; (b) Scatter plot of N ratiot; (c) Line plot of GC skew; (d) Heatmap of gene density; (e) Bar plot of GC ratio.

### Mitochondrial genome assembly

The mitochondrial genome was assembled using Illumina data with MitoZ version 3.6^19^, which was further employed for mitochondrial annotation. To ensure accuracy, the assembled mitochondrial genome was compared against the nuclear genome using BLAST+ version 2.16.0^20^, and the verified mitochondrial sequence was incorporated into the final genome assembly. Notably, *Meretrix* sp. MF1 and *Meretrix lamarckii* JML1 exhibited the closest match to the same species, *Meretrix lamarckii*, albeit from distinct sources. Specifically, *Meretrix* sp. MF1 showed the highest similarity to Sequence ID: NC_016174.1 (GenBank), while *Meretrix lamarckii* JML1 showed the highest similarity to Sequence ID: KP244451.1. Furthermore, mitochondrial data revealed an additional tRNA-Leu in *Meretrix* sp. MF1 compared to *Meretrix lamarckii* JML1, potentially indicating distinct species status. In addition, *Meretrix* sp. MT1 was found to be most closely related to *Meretrix lusoria* (Sequence ID: NC_014809.1). A summary of all assembled mitochondrial data is provided in Table 4.

**Table 4.** Summary statistics of the *Meretrix* mitochondrial genome.

| Species | *Meretrix* sp. MF1 | *Meretrix* sp. MT1 | *M. lamarckii* JML1 |
| --- | --- | --- | --- |
| Length (bp) | 20,025 | 19,263 | 19,919 |
| Circularity | Yes | Yes | Yes |
| Closely related species (from NCBI) | *M. lamarckii* (NC_016174.1) | *M. lusoria* (NC_014809.1) | *M. lamarckii* (KP244451.1) |
| Protein coding genes | 13 | 13 | 13 |
| tRNA genes | 23 | 22 | 22 |
| rRNA genes | 2 | 2 | 2 |
| Genes totally found | 38 | 37 | 37 |

### Repetitive sequence identification

RepeatModeler version 2.0.5^21^ and RepeatMasker version 4.1.5^22^ were used to analyze the *Meretrix* genome assemblies, enabling the *de novo* identification of transposable elements (TEs) and the classification of repetitive and low-complexity sequences (Table 5). The total proportion of repetitive elements in *Meretrix* sp. MF1, *Meretrix* sp. MT1, and *M. lamarckii* JML1 genomes were 41.57%, 41.75%, and 40.35%, respectively, with unclassified repeats accounting for 32.30%, 32.40%, and 31.36%. In terms of TE composition, Retroelements (Class I) were identified, constituting 6.99%, 6.55% and 6.64% of the genomes, respectively. The DNA transposons (Class II) were 1.98%, 1.59% and 1.77%, respectively. The consistent repeat content and distribution patterns across the three *Meretrix* species suggest a conserved genome organization and repetitive element dynamics within the genus.

**Table 5.**
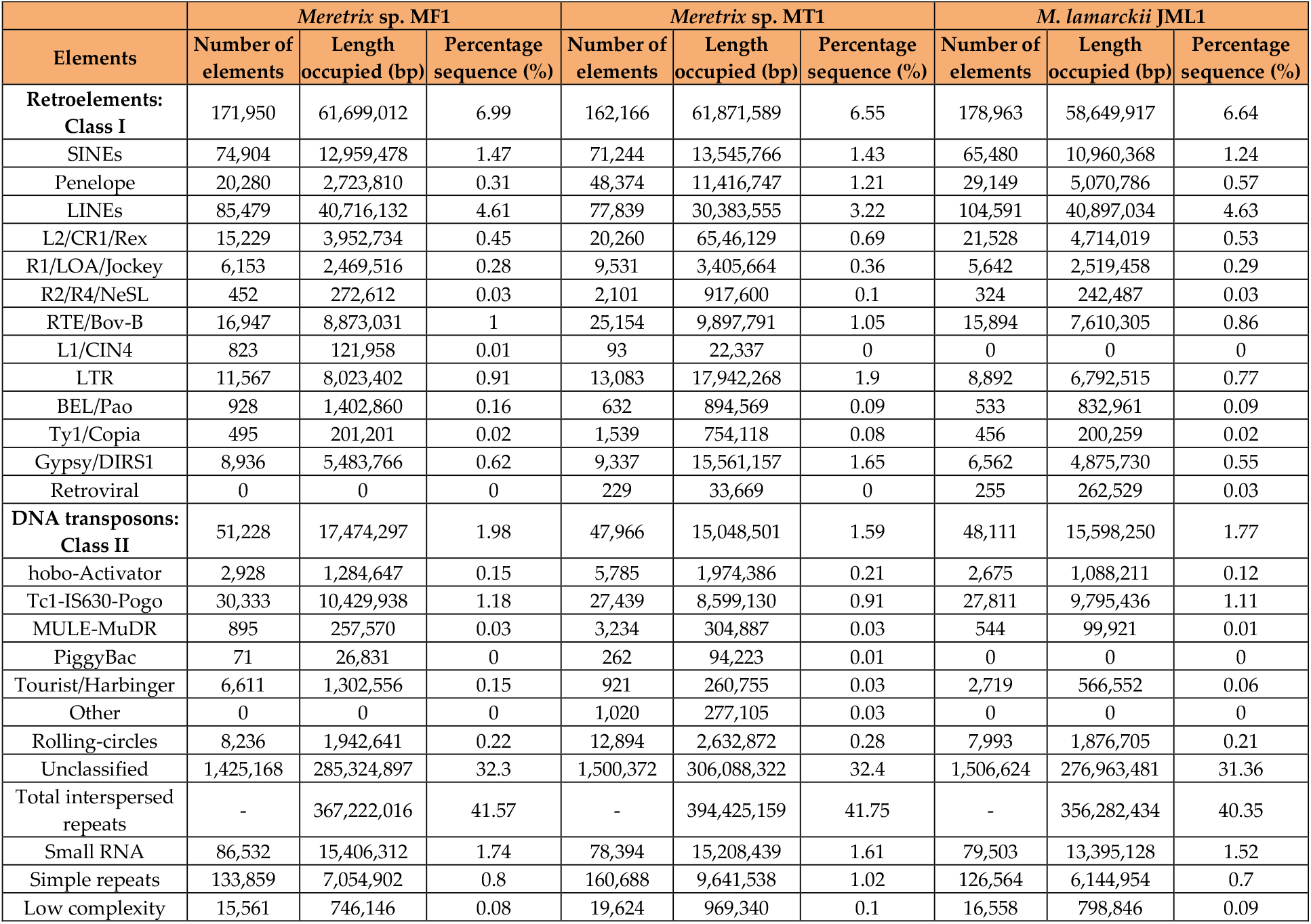
Repetitive Element Composition of the *Meretrix* Genome Assembly.

### Gene prediction and functional annotation

Gene prediction was performed on a genome version that was soft-masked for repeats using RepeatMasker version 4.1.5^22^. The prediction was carried out with BRAKER version 3.0.8^23^, employing a protein evidence-based approach using Metazoa dataset from OrthoDB version 12^24^. Gene prediction for *Meretrix* sp. MF1 was performed using BRAKER, which initially predicted 45,263 genes and 49,050 transcripts. Subsequently, the selectSupportedSubsets.py script within BRAKER was employed to filter transcripts based on hint support, resulting in a subset of 32,329 transcripts. Transposable elements (TEs) were then masked using TEsorter version 1.2.7^25^, yielding a final set of 30,417 transcripts. Functional annotation was conducted using EggNOG-mapper version 2.1.12^26^ and InterProScan version 5.73-104.0^27,28^, to identify protein homologs, which included six database resources: eggNOG, Gene Ontology (GO) terms, Kyoto Encyclopedia of Genes and Genomes (KEGG), InterPro, Protein ANalysis THrough Evolutionary Relationships (PANTHER), and Pfam. A total of 25,531 genes were successfully annotated with functional information from at least one of these databases. Comprehensive gene annotation statistics for the *Meretrix* genome are provided in Supplementary Table 1.

### Genomic similarity comparison and evolutionary analysis

FastANI version 1.34^29^ was applied to calculate the ANI among the genomes of *Meretrix* sp. MF1, *Meretrix* sp. MT1, *M. lamarckii* JML1, and *M. petechialis*. The results revealed that the ANI between *Meretrix* sp. MF1 and *M. lamarckii* JML1 was 94.33% (other comparisons are provided in Supplementary Table 2), suggesting that *Meretrix* sp. MF1 might represent a potentially novel species in Taiwan. We propose the name *M. formosana*. To explore evolutionary relationships, BUSCO version 5.7.1^30^ was used to extract conserved Metazoa homologous genes from 11 genomes of Veneridae, including *Callista chione, Cyclina sinensis, Mercenaria mercenaria, M. lamarckii* JML1, *M. petechialis, Meretrix* sp. MF1, *Meretrix* sp. MT1, *Mysia undata, Ruditapes philippinarum, Saxidomus purpurata*, and *Venus verrucosa* (Supplementary Table 3). Multiple sequence alignment was performed using MUSCLE version 5.3^31^, followed by trimming with trimAI version 1.5.0^32^ to generate the supermatrix alignment file. A phylogenetic tree was constructed based on the concatenated alignments using IQ-TREE version 1.6.12^33^, incorporating divergence times estimates obtained from the TimeTree database^34^ (accessed on Feb. 10, 2025). The estimated divergence times included 194 million years between *M. mercenaria* and *V. verrucosa*, 171 million years between *V. verruco*sa and *R. philippinarum*. The final phylogenetic tree was visualized using MEGA version 11.0.13^7^, with *M. mercenaria* as the outgroup (Fig. 3).

**Figure 3.**
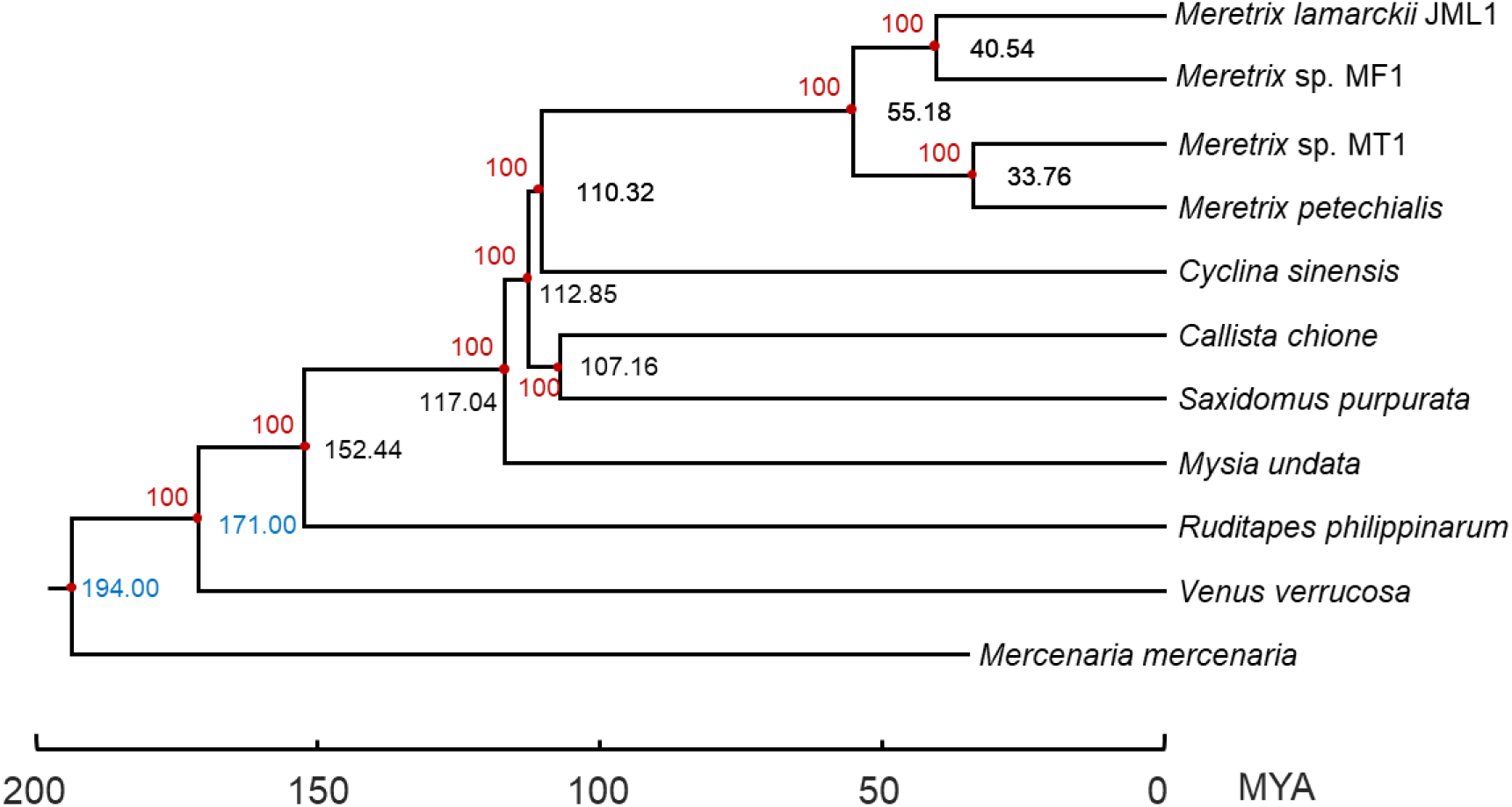
Estimated divergence times among *Meretrix* species inferred from metazoan orthologous genes. Phylogenetic tree of 11 mollusk species, rooted with *Mercenaria mercenaria* as the outgroup. Bootstrap values are shown in red next to each node. Divergence time estimates from the TimeTree database are indicated by blue. Estimated divergence times between species pairs are listed next to each node. Mya: million years ago.

## Data Records

All raw sequencing data have been deposited in the BioProject at NCBI under accession number PRJNA1227740^35^.

The Illumina data were deposited in the Sequence Read Archive at NCBI under accession number SRR32575144, SRR32575146, and SRR32575149^36-38^.

The Nanopore data were deposited in the Sequence Read Archive at NCBI under accession number SRR32575145, SRR32575147, and SRR32575150^39-41^.

The Hi-C data were deposited in the Sequence Read Archive at NCBI under accession number SRR32575148^42^.

## Technical Validation

### Genome assembly and annotation completeness evaluation

To assess the completeness and accuracy of the assembled genomes, multiple quality assessment tools were utilized. First, BUSCO version 5.7.1^30^ with the metazoa_odb10 lineage database, was used to evaluate the genome completeness. In the *Meretrix* sp. MF1 genome, 910 (95.4%) single-copy ortholog were fully identified, while *Meretrix* sp. MT1 and *M. lamarckii* JML1 contained a complete set of 884 (92.7%) and 874 (91.6%) single-copy orthologs, respectively. The completeness scores for all three species exceeded 92.9% based on both the eukaryote_odb10 and metazoa_odb10 database, demonstrating the high quality and completeness of the assembled genomes (Table 6). Next, Merqury version 1.3^43^ was used to evaluate genome completeness using a *k*-mer-based approach. *K*-mers derived from Nanopore data were analyzed to calculate the quality value (QV) score, resulting in 97.62% *k*-mer completeness and an assembly consensus QV of 49.74 in *Meretrix* sp. MF1 (Supplementary Table 4). The statistical results for *Meretrix* sp. MT1 and *M. lamarckii* JML1 are also presented in Supplementary Table 4. To further assess assembly accuracy, Illumina reads were aligned to the genome using BWA version 0.7.18^44^. Statistical analysis with SAMtools version 1.21^45^ showed that 99.72% of the Illumina reads successfully mapped to the genome, achieving a coverage of 98.25%, confirming the high accuracy of the assembly (Supplementary Table 5). The results for *Meretrix* sp. MT1 and *M. lamarckii* JML1 are also presented in Supplementary Table 5. Additionally, Juicebox version 1.11.08^46^ was employed to visualize the assembled scaffolds and detect potential misassemblies. Manual inspection revealed no characteristic patterns of read coverage indicative of misjoins, translocations, or inversions.

**Table 6.** Results of BUSCO completeness assessment for the Meretrix genome assembly.

| Species | *Meretrix* sp. MF1 |  | *Meretrix* sp. MT1 |  | *M. lamarckii* JML1 |  |
| --- | --- | --- | --- | --- | --- | --- |
|  | eukaryota_odb10 | metazoa_odb10 | eukaryota_odb10 | metazoa_odb10 | eukaryota_odb10 | metazoa_odb10 |
| Complete BUSCOs (C) | 251 (98.4%) | 918 (96.2%) | 244 (95.7%) | 889 (93.2%) | 242 (94.9%) | 886 (92.9%) |
| Complete and single-copy BUSCOs (S) | 249 (97.6%) | 910 (95.4%) | 243 (95.3%) | 884 (92.7%) | 242 (94.9%) | 874 (91.6%) |
| Complete and duplicated BUSCOs (D) | 2 (0.8%) | 8 (0.8%) | 1 (0.4%) | 5 (0.5%) | 0 (0.0%) | 12 (1.3%) |
| Fragmented BUSCOs (F) | 0 (0.0%) | 7 (0.7%) | 1 (0.4%) | 13 (1.4%) | 6 (2.4%) | 22 (2.3%) |
| Missing BUSCOs (M) | 4 (1.6%) | 29 (3.1%) | 10 (3.9%) | 52 (5.4%) | 7 (2.7%) | 46 (4.8%) |
| Total BUSCO groups searched | 255 | 954 | 255 | 954 | 255 | 954 |

## Code availability

### Genome annotation

1. RepeatModeler: parameters: all parameters were set as default.
2. RepeatMasker: parameters: -e rmblast -lib database_repeat-families.fa genome.fasta -xsmall -s -gff.
3. Braker3: parameters: --genome=genome.fa --prot_seq=proteins.fa -- gff3.

### Genome assembly

1. NextDenovo: parameters: all parameters were set as default.
2. Masurca: parameters: all parameters were set as default.
3. NextPolish: parameters: rerun = 2. sgs_options = -max_depth 100 -bwa. lgs_minimap2_options = -x map-ont.

### Orthologous genes analysis

1. BUSCO: parameters: -i genome.fa -r -o Busco_result --auto-lineage- -m geno -f --offline –augustus.
2. iqtree: parameters: iqtree -s SUPERMATRIX -m TEST -bb 1000 -alrt 1000.

## Acknowledgements

This study was supported by the National Science and Technology Council of Taiwan (MOST 111-2628-M-019-001-MY3, and 113-2119-M-001-011-).

## Author contributions statement

Y.N.H. conceived and supervised the study. C.C.C., H.Y.L., T.H.H., and Y.N.H. collected the sample. C.C.C. performed the laboratory work. C.C.C and Y.N.H. performed bioinformatics analysis. C.C.C. and H.Y.L. drafted the manuscript. T.H.H., S.L.T., and Y.N.H. provided review and modification of the manuscript. All authors read and approved of the final manuscript.

## Competing Interests

The authors declare no competing interests.

## Notes

https://doi.org/10.6084/m9.figshare.28674617.v2

### Competing Interest Statement

The authors have declared no competing interest.

